# Detection of *Echinococcus* spp. and other taeniid species in lettuces and berries: two international multicenter studies from the MEmE project

**DOI:** 10.1101/2024.06.10.598207

**Authors:** Gérald Umhang, Fanny Bastien, Alexandra Cartet, Haroon Ahmad, Kees van der Ark, Rebecca Berg, Piero Bonelli, Rebecca K. Davidson, Peter Deplazes, Gunita Deksne, Maria João Gargate, Joke Van der Giessen, Naila Jamil, Pikka Jokelainen, Jacek Karamon, Selim M’Rad, Pavlo Maksimov, Myriam Oudni-M’Rad, Gillian Muchaamba, Antti Oksanen, Paola Pepe, Marie-Lazarine Poulle, Laura Rinaldi, Małgorzata Samorek-Pieróg, Federica Santolamazza, Azzurra Santoro, Cinzia Santucciu, Urmas Saarma, Manuela Schnyder, Isabelle Villena, Marion Wassermann, Adriano Casulli, Franck Boué

## Abstract

Cystic and alveolar echinococcosis are severe zoonotic diseases characterized by long asymptomatic periods lasting months or years. Viable *Echinococcus* eggs released into the environment through the feces of canids can infect humans through accidental ingestion via hand-to-mouth contact or consumption of contaminated food or water. Both *Echinococcus multilocularis* and *Echinococcus granulosus sensu lato* are considered as foodborne parasites. However, when considering possible pathways of human infection, it appears that food and water-borne related variables do not significantly increase the risk of infection. Providing evidence-based data for the presence of DNA and, potentially, eggs in fresh produce is crucial in understanding foodborne transmission of *Echinococcus* spp. to humans. Two multicenter and multicountry studies were conducted within the One Health EJP framework to estimate the proportion of lettuces and berries contaminated by *E. multilocularis*, *E. granulosus s.l*., and other taeniid DNAs from 12 European countries, Tunisia and Pakistan. A total of 1,117 lettuces, 300 strawberries and 130 blueberries samples were collected and analyzed by washing, sequential sieving and real-time PCRs. *Echinococcus multilocularis* DNA was detected in 1.2% (7/570) of samples tested from the seven European endemic areas and in 2% (2/100) from Pakistan. In the five European endemic countries for *E. granulosus s.l.*, *E. granulosus sensu stricto* DNA was identified in 1.3% of lettuces (9/695). The proportion of lettuces contaminated by *E. granulosus sensu stricto* DNA was 12% (9/75) in Tunisia and 4% (4/100) in Pakistan. Regarding berries, *E. multilocularis* DNA was detected in 5.4% of strawberries (n=11/202), 7.3% of blueberries (6/82) from the seven European endemic countries and 56% of blueberries (14/25) from Pakistan. High contamination rates of *E. granulosus s.s* were found outside of Europe, with 12.0% in blueberries (3/25) from Pakistan and 81.3%. DNA in strawberries (13/16) from Tunisia. The total contamination rate of all taeniid species DNA in lettuces (5.3%; 59/1,117) and berries (12.1%; 58/480) suggests that the transfer of taeniid eggs from carnivore feces to food is not uncommon. Although we assume that eggs are the source of the DNA detected in this study, the viability of such eggs is unknown. The detection of *Echinococcus* species in lettuces and berries suggests a potential risk of foodborne human infection. The relative contribution of this risk remains to be estimated. Further studies on food and environmental contamination are necessary to cover different epidemiological contexts and social habits, leading to a better understanding of human infections by *Echinococcus* eggs.

## 1. Introduction

The echinococcosis disease group is listed among the Neglected Tropical Diseases (NTDs) of global public health importance prioritized by the World Health Organization (WHO) for their control (WHO, 2024; Casulli and Tamarozzi, 2021). Both cystic (CE) and alveolar (AE) echinococcosis are characterized by a long asymptomatic period lasting months or years. CE is a chronic and disabling disease characterized by the development of parasitic cysts, mainly located in the liver (70%), lungs (19%) or in uncommon localizations (11%) as bones, hearth, muscles and central nervous system (Casulli, Pane, et al., 2023). The concentric growth of the cysts may have an effect of compression on neighboring organs and tissues, which sometimes can compromise their function. The mortality rate for CE is estimated around 2–4% but can be higher if medical treatment and care are inadequate (Brunetti et al., 2010). An estimate of around 65,000 human cases of CE was reported from 1997 to 2021 in 40 selected European countries (Casulli, Abela-Ridder, et al., 2023). AE is generally considered to be a more serious life-threatening disease than CE, as it is characterized by a tumor-like infiltrative parasitic tissue (vesicles), mainly present in the liver with parasitic metastasis spreading in the neighboring organs (Gottstein et al., 2017). AE is distributed in the Northern Hemisphere, while CE is present worldwide. According to non-updated estimates, the annual global incidence of AE was 18,000 cases per year in 2010, of which 91% occurring in China (Torgerson et al., 2010). During the period 2018-2022, official statistics from the 27 member states of the European Union have reported an average of 690 unspecified echinococcosis human cases per year, including 338 CE and 151 AE human cases (EFSA, 2022). Nevertheless, it should be kept in mind that considerable underreporting is existing for both diseases with a heterogeneity in data reporting among European countries (van der Giessen et al., 2021).

*Echinococcus multilocularis* and *Echinococcus granulosus sensu lato* (*s.l*.) are the causative agents of AE and CE, respectively. The lifecycle of these two parasites is based on the trophic relationship between canids as definitive hosts and ungulates or rodents as intermediate hosts. In non-Arctic areas of Europe, the red fox (*Vulpes vulpes*) is the primary definitive host for *E. multilocularis*, while in Asia, the dog is also heavily involved due to their frequent predation on small rodents (Romig et al., 2017). The life cycle of *E. granulosus s.l*., which is a complex of cryptic species, is primarily domestic. The dog serves as the main definitive host, with various livestock species acting as intermediate hosts, depending on the specific *E. granulosus* species involved. *E. granulosus s.l*. is currently represented by five species (genotypes), namely *E. granulosus sensu stricto* (*s.s*.; G1 and G3), *E. equinus* (G4), *E. ortleppi* (G5), *E. canadensis* (G6/7 complex, G8 and G10) and *E. felidis* (Romig et al., 2017; Vuitton et al., 2020). The latter species has never been documented to infect humans and was only reported from Africa. The other four species are present in Europe and worldwide with varying frequencies and geographical distributions in humans (Alvarez Rojas et al., 2014; Casulli et al., 2022). Compared to *E. granulosus s.l*., the genetic diversity of *E. multilocularis* is considered relatively low (Haag et al., 1997; Knapp et al., 2015). Four distinct clades of *E. multilocularis* are currently recognized based on their geographical origin, as initially determined by mitochondrial sequencing (Nakao et al., 2009) and further supported by EmsB nuclear microsatellite analysis (Knapp et al., 2007; Umhang et al., 2021). Nevertheless, Asian-like samples have been reported in Europe and European-like samples in North America (Karamon et al., 2017; Santa et al., 2023).

Viable *Echinococcus* eggs can infect humans through accidental ingestion via hand-to-mouth contact or consumption of contaminated food or water. The eggs are released into the environment through the feces of canids (Alvarez Rojas et al., 2018; Kapel et al., 2006) and are highly resistant to temperature variations under natural conditions. Eggs of *E. multilocularis* can survive up to 240 days in autumn-winter and 78 days in summer, as reported in Germany (Veit et al., 1995). Similarly, eggs of *E. granulosus s.l.* can survive up to 41 months, as documented in Argentinian Patagonia (Sánchez Thevenet et al., 2019). Furthermore, taeniid eggs can be dispersed passively over short and long distances by wind, rain, insects, or birds (Benelli et al., 2021; Raymond and St Clair, 2023; Sánchez Thevenet et al., 2019), leading to contamination of various environmental matrices. However, there is limited information available on environmental contamination caused by these parasites in different matrices, such as fox or dog fur, soil, water, vegetables, and fomites (Tamarozzi et al., 2020).

*Echinococcus multilocularis* and *E. granulosus s.l.* are both considered foodborne parasites, ranking first and fourth in Europe and second and third in the world, respectively (Bouwknegt et al., 2018; Devleesschauwer et al., 2017). Two systematic reviews and meta-analyses were conducted on AE and CE using previously published case-control and cross-sectional studies to identify potential risk factors (Conraths et al., 2017; Possenti et al., 2016). The results of both studies suggested that, when considering possible pathways of human infection, food- and water-borne related variables do not significantly increase the risk of infection with *Echinococcus* spp. Another systematic review and meta-analysis from Torgerson et al. (2020), used attributable fractions for the potential sources of CE and AE infections, instead of considering risk factors. This study confirmed that food can be a source of human infections for CE worldwide. This association was not statistically significant in regions with a high incidence of AE, such as China, whereas it was significant in regions with a low incidence, such as Europe (Torgerson et al., 2020). Only a few recent studies have provided data on the detection of *E. multilocularis* DNA in raw vegetables, berries, and mushrooms. Large differences in detection rates have been observed, ranging from 1.3% in lettuces (n=157) purchased in Switzerland to 23.3% in forest fruits, vegetables, and mushrooms (n=103) collected in Poland (Guggisberg et al., 2020; Lass et al., 2015a). Regarding *E. granulosus s.l.*, M’Rad et al. (2020) found that 1.3% of vegetables (n=240), purchased from local markets in Tunisia, were contaminated with *E. granulosus s.s*. eggs. Comparing the results between these studies is challenging due to the use of different methods (for the enrichment of samples, DNA extraction and molecular detection), with generally unevaluated performance or high detection limits of these tests. Providing evidence-based data for the presence of DNA and potentially eggs in fresh products is crucial in understanding foodborne transmission of *Echinococcus* spp. to humans (Bouwknegt et al., 2018). This task is challenging since it requires a method to enrich matrices and detect very low number of taeniid eggs in food items, combined with molecular tests to identify the species involved. In this context, multicenter and large-scale studies, utilizing the same protocols for the detection of the parasites is mandatory. Therefore, we took advantage of the MEmE project (Multi-centre study on *Echinococcus multilocularis* and *Echinococcus granulosus s.l.* in Europe: development and harmonisation of diagnostic methods in the food chain) within the One Health EJP framework to conduct a multicenter and multicountry studies. The aim of this study was to estimate the proportion of lettuces and berries contaminated by *E. multilocularis*, *E. granulosus s.l*., and others taeniid species DNAs on a large geographical scale.

## 2. Materials and methods

### 2.1. Evaluation of the limit of detection and robustness of the method

A preliminary validation step of the method was conducted to confirm its sensitivity from samples artificially spiked with *E. multilocularis* eggs. The method used in this study, to detect DNA from taeniid eggs in both lettuces and berries, was previously described by Guggisberg et al. (2020). The method involved three steps: 1) washing the samples, 2) sequential sieving of the washing solution to obtain a pellet from the 20µm filter, and 3) DNA extraction (using here DNeasy Blood & Tissue kit, Qiagen) from the pellet for molecular analysis to detect *E. multilocularis* using a real-time PCR (Knapp et al., 2016). Lettuce heads and strawberry trays were purchased from supermarkets. These two matrices, comprising 300g of lettuce leaves or 200g of strawberries, respectively, were spiked with a defined number of *E. multilocularis* eggs before being washed, filtered, and molecularly identified. To validate the process, a preliminary step consisting of spiking 10 and 5 lettuce samples with 10 and 5 eggs, respectively, was conducted. For strawberries, 3 samples were spiked with 10 eggs and 2 samples with 5 eggs. Subsequently, 3 replicates composed by 24 lettuce samples each and 3 replicates of 8 strawberry samples each, were spiked with 3, 2 or 1 eggs each sample. The limit of detection (LOD) was defined as the minimum number of eggs that can be detected in at least 23 out of the 24 replicates for lettuces or all 8 replicates for strawberries. All eggs used for spiking were obtained from the feces of experimentally infected red foxes, as approved by the Anses/ENVA/UPEC ethics committee and the French Ministry of Research (Apafis n°33541-202110081648536) and inactivated at −80°C before use.

In summer 2020, a second preliminary validation step was carried out by sampling lettuces collected from the field (i.e. local markets and private kitchen gardens) in north-eastern France to test the robustness of the approach, before the implementation of the method in other countries.

### 2.2. Sampling of lettuces and berries

The first multicenter study focused on lettuces, with sampling conducted during summer 2021 and involving 15 laboratories from 12 countries: Denmark (n=1), France (n=1), Germany (n=2), Italy (n=3), Latvia (n=1), the Netherlands (n=1), Norway (n=1), Pakistan (n=1), Poland (n=1), Portugal (n=1), Switzerland (n=1) and Tunisia (n=1). Each laboratory was asked to collect between 50 and 100 heads of lettuce. Such collection should have been carried out in local markets (4 samples from the same site), private kitchen gardens (2 from the same site) or from supermarkets (4 from each shop). Additionally, six laboratories (from Campania and Sardinia regions of Italy, Latvia, north of Germany and Switzerland) collected other green leafy vegetables, mainly chard and parsley.

In summer 2022, a second multicenter study was organized, focusing on berries (mainly strawberries and blueberries). In this study, blueberries will refer to low-growing Eurasian blueberries (also named bilberries and corresponding to *Vaccinium angustifolium*, *V. boreale*, *V. mytilloides* and *V. pallidum*) and not to North American blueberries (*V. darrowii* and *V. corymbosum*) growing on taller, cultivated bushes. Twelve laboratories from 12 countries were involved in this study on berries: Denmark, Estonia, Finland, France, Italy, Latvia, the Netherlands, Pakistan, Poland, Portugal, Switzerland and Tunisia. Each laboratory was asked to collect between 20 and 30 berry samples (100-200 g each). Each laboratory collected berry samples from local markets, directly in the field and forest or with supplementation from supermarkets. Only one sample per site was requested. The European countries were considered to be endemic for *E. granulosus* or *E. multilocularis* according to EFSA and ECDC (EFSA, 2022).

### 2.3. Analysis of field samples for the detection of *E. multilocularis*, *E. granulosus s.l.* and others taeniid species

The samples collected from the field in the different countries were processed as following: 1) washing the samples with subsequent sedimentation of the rinsing liquid to obtain a pellet, 2) sequential sieving of the pellet to obtain an enriched sediment from the last 20µm filter, and 3) DNA extraction and molecular detection of the enriched pellet. All participants conducted step 1 (washing and sedimentation) to ensure a prompt analysis of fresh products within a few days of purchase, thus avoiding any decomposition of the sample. Standard operating procedures for the first step of washing and sedimentation of lettuces and berries were distributed to each participant prior to sampling, to ensure a reproducibility of the method. The participants in the two multicenter studies were instructed not to wash the food items before starting the analysis (step 1). Steps 2 and 3, which involved filtration, DNA extraction, and molecular detection, were carried out in a single laboratory (Anses LRFSN, Nancy, France) to ensure reproducibility of the method and sensitive detection of parasite DNA. In Step 1, the washing process consisted of mixing 500 ml of 0.02% Tween with up to 300 g of lettuce leaves or up to 250 g of berries. This initial washing step used standard plastic bags (Guggisberg et al., 2020) provided by Anses LRFSN (Nancy, France) to avoid potential bias with the use of different materials. External lettuce leaves were not removed from the sample and were included in the washing process. For berries, the vigorous manual washing step from the original protocol for lettuce was replaced by an orbital shaker (2 sessions of 15 minutes each) to remove as much residue as possible from the berries. The berries were also manually mixed between the two shaking sessions to ensure that the entire surface of each berry was washed. After washing, the rinsing liquid solution was transferred to another plastic bag for overnight sedimentation to obtain a pellet. The pellet was stored at +4°C or −20°C before being transferred to France (Anses, Nancy laboratory) for steps 2 and 3. All lettuce and berry samples collected in France were processed continuously without the sedimentation step after washing that was necessary only for the other partners.

During step 2, pools were created for sequential sieving by grouping two sediment samples together if they originated from the same site. Most of the lettuce samples (93.5%) and a third of berry samples (34.4%) were analyzed by pooling samples. If a positive molecular test was obtained from a pooled samples, it was assumed that only one of the two samples was positive when estimating the proportion of contamination. The samples were filtered sequentially through 105 µm, 40 µm, and 20 µm mesh size. The final pellet deposited on the last 20 µm mesh filter was rinsed with 0.02% Tween using a pipette.

In Step 3, following brief centrifugation to remove the supernatant, DNA was extracted from the pellet using the conventional DNeasy Blood & Tissue kit (Qiagen) with an initial overnight lysis step to optimize recovery of DNA from eggs. The real-time PCR method described by Knapp et al. (2016) was used to detect *E. multilocularis* DNA. *Echinococcus granulosus s.l.* DNA was detected using primers and probes for the specific detection of *E. granulosus s.s*., *E. canadensis* and *E. ortleppi* (Maksimov et al., 2020). Others taeniid species were identified through conventional PCR (Trachsel et al., 2007) and confirmed by Sanger sequencing performed by a private company (Eurofins GmbH, Germany).

### 2.4. Statistics

The detection rates of the different parasite species for each type of matrix or sources of supply (kitchen garden, local markets, supermarkets or in the field or forest) were expressed as a percentage with 95% confidence intervals. Comparisons based on matrix type were conducted using either a chi-square test or a Fischer test if the sample size was too small.

## 3. Results

### 3.1. Estimation of the limit of detection

Based on the preliminary validation, the LOD for lettuces was estimated to be 3 eggs, as *E. multilocularis* DNA was detected in 23 out of 24 lettuce head replicates (detection rate >95%) (Supplementary Table 1). The detection rate of lettuce replicates spiked with 2 and 1 eggs was 75% (18/24) and 50% (12/24), respectively (Supplementary Table 1). The LOD for *E. multilocularis* DNA in strawberry replicates was estimated to be 3 eggs based on the detection of *E. multilocularis* DNA in all 8 samples spiked with 3 eggs. The detection rate for both 2 and 1 spiked eggs was 88% (7/8) (Supplementary Table 1).

### 3.2. Detection of DNA of *E. granulosus s.l., E. multilocularis* and other taeniid species in lettuces and other leafy vegetables

Out of the 1,117 collected lettuces, 570 were from *E. multilocularis* endemic areas of European countries (Denmark, France, Germany, Latvia, the Netherlands, Poland and Switzerland) (Table 1). *Echinococcus multilocularis* DNA was detected in 7 lettuce samples (1.2%; 7/570) from four European countries (France, Denmark, Latvia and Switzerland) (Table 1). The proportions of detection of *E. multilocularis* DNA in lettuces was comparable (around 1%) for most of the countries where *E. multilocularis* was detected: Denmark (4%; 2/50), France (1.3%; 3/226), Latvia (1.6%; 1/62) and Switzerland (1.3%; 1/80). Two samples (2%) out of 100 from Pakistan tested positive for *E. multilocularis* DNA. Furthermore, among the 55 non-lettuce vegetable samples collected from endemic areas, one *E. multilocularis* positive sample (1.8%; 1/55) was found in a chard from northern Germany (Supplementary Table 2).

**Table 1:** Number of tested positive samples and detection proportions of *Echinococcus multilocularis, Echinococcus granulosus sensu lato* and others taeniid species DNA in lettuces from the different countries. The 95% confidence intervals are indicated between brackets. *Proportions were calculated considering only those European countries for which areas endemic for *E. multilocularis* have been sampled (Denmark, France, Germany, Latvia, the Netherlands, Poland and Switzerland) and European countries where a domestic lifecycle is established for *E. granulosus* (France, Italy, Latvia, Poland, Portugal).

| Country (regions) | number of strawberries samples | *E. multilocularis* |  | *E. granulosus s.l.* |  | Other taeniid species |  |
| --- | --- | --- | --- | --- | --- | --- | --- |
|  |  | number of samples tested positive with *E. multilocularis* | proportions of positive cases (95% CI) | number of samples tested positive with *E. granulosus s.l.* | proportions of positive cases (95% CI) | number of samples tested positive with others taeniid species | proportions of positive cases (95% CI) with others taeniid species |
| Denmark | 56 | 1 | 1.8% (0-9.6) | 0 | 0% (0-63.8) | 1 *Hydatigera* sp. | 1.8% (0-9.6) |
| Estonia | 30 | 5 | 16.7% (5.6-34.7) | 0 | 0% (0-52.2) | 0 | 0% (0-52.2) |
| Finland | 10 | 0 | 0% (0-30.8) | 0 | 0% (0-30.8) | 0 | 0% (0-30.8) |
| France | 54 | 0 | 0% (0-6.6) | 0 | 0% (0-6.6) | 0 | 0% (0-6.6) |
| Italy (Sardinia) | 55 | 0 | 0% (0-6.5) | 1 | 1.8% (0-9.7) | 2 *Hydatigera* sp. | 3.6% (0.4-12.5) |
| Latvia | 30 | 4 | 13.3% (3.8-30.7) | 0 | 0% (0-11.6) | 0 | 0% (0-11.6) |
| Netherlands | 8 | 1 | 12.5% (0.3-52.7) | 0 | 0% (0-36.9) | 0 | 0% (0-36.9) |
| Poland | 13 | 0 | 0% (0-24.7) | 1 | 7.7% (0.2-36.0) | 0 | 0% (0-24.7) |
| Portugal | 17 | 0 | 0% (0-19.5) | 1 | 5.9% (0.1-28.9) | 0 | 0% (0-19.5) |
| Switzerland | 11 | 0 | 0% (0-28.5) | 0 | 0% (0-28.5) | 0 | 0% (0-28.5) |
| **SubTotal Europe\*** | 284 | 11 | 5.4%* (2.7-9.5) | 3 | 1.5%* (0.3-4.3) | 3 *Hydatigera* sp. | 1.0 (0.2-3.0) |
| Tunisia | 16 | 0 | 0% (0-20.6) | 13 | 81.3% (54.4-96.0) | 1 *T. hydatigena* | 6.3% (0.2-30.2) |
| **Total** | 300 | 11 | 3.7% (1.8-6.5) | 16 | 5.3% (3.1-8.5) | 3 *Hydatigera* sp.,<br>1 *T. hydatigena* | 1.3% (0.4-3.3) |

The overall contamination rate of *E. granulosus s.l.* among lettuce samples was 2% (22/1,117) (Table 1). The contamination rate, if we only consider European countries, was 1% (9/942), with variations between areas ranging from 0 to 6.5%. Except for one case of *E. canadensis* species detected in Pakistan, the other findings were all *E. granulosus s.s*. An average rate of 3.4% (8/232) was obtained for the three Italian regions (Abruzzo, Campania and Sardinia) and 1.6% (1/62) in Latvia. In Tunisia, the rate of lettuce contamination was the highest at 12% (9/75), while in Pakistan, the rate was 4% (4/100). No positive sample of *E. granulosus s.l.* were detected in leafy vegetables other than lettuce (Supplementary Table 2).

The total proportion of other taeniid species detected in lettuces was 2.5% (28/1,117) (Table 1). In Europe, the proportion of 1.7% (16/942) was lower compared to Tunisia (4.0%; 3/75) and Pakistan (9.0%; 9/100). Fifteen samples of *Hydatigera* sp. (1.3%; 15/1,117), six of *Taenia hydatigena* (0.5%; 6/1,117), and five of *Taenia saginata* (0.4%; 5/1,117), were identified, as well as one of *Taenia multiceps* (0.1%). One of the three Tunisian lettuce samples with *T. hydatigena*, also tested positive for *E. granulosus s.s*. In addition, the partial 12S rRNA gene sequence obtained from a sample of lettuce from Norway did not allow differentiation between *Taenia serialis* and *Taenia krabbei*. *Hydatigera* sp. was also detected in chard, basil, and sorrel samples from Latvia and Norway (Supplementary Table 2). The DNA of the cestode species *Atriotaenia incisa* was identified in one lettuce sample from Latvia, as the Cest4-5 primers have a wider amplification spectrum than just taeniid species (Trachsel et al., 2007).

### 3.3. Detection of DNA of *E. granulosus s.l.*, *E. multilocularis* and other taeniid species in berries

A total of 480 berry samples were collected, of which 300 were strawberries, 130 blueberries and the remaining 50 samples were from eight other berry species: blackberries (n=13), raspberries (n=21), red currants (n=5), blueberry bushes (n=4), red lingonberries (n=3), black currants (n=2), white currants (n=1), Saskatoon berries (n=1). A total of 202 strawberry samples and 82 blueberry samples were collected from European countries where *E. multilocularis* is endemic (Denmark, Estonia, France, Latvia, the Netherlands, Poland, and Switzerland). *Echinococcus multilocularis* DNA was detected in 5.4% (11/202) of the strawberry samples (Table 2) and 7.3% (6/82) of the blueberry samples (Table 3) from these endemic European countries. No *E. multilocularis* DNA was detected in strawberry or blueberry samples from Poland or Switzerland. In the other European endemic countries, the level of contamination ranged from 1.8% to 16.7%. The two Baltic countries had a high proportion of contamination, with 13.3% (4/30) in both strawberries and blueberries from Latvia and 16.7% (5/30) in strawberries from Estonia. Although no positive strawberry sample (0/54) was found in France, *E. multilocularis* DNA was identified in 3.2% (1/31) of blueberries analysed. *E. multilocularis* DNA was also detected in The Netherlands from blueberries (1/6) and strawberries (1/8) at 16.7% and 12.5%, respectively. In Denmark, *E. multilocularis* DNA was detected in 1.8% (1/56) of strawberry samples. A high proportion (56%; 14/25) of blueberries from Pakistan were also contaminated with *E. multilocularis* DNA. *Echinococcus multilocularis* DNA was not found in other types of berries (Supplementary Table 3).

**Table 2:** Number of cases and proportions of detection of *E. multilocularis, E. granulosus sensu lato* and other taeniid species DNA in stawberries from the different countries. The 95% confidence intervals are indicated between brackets. *Proportions were calculated considering only those European countries for which areas endemic for *E. multilocularis* have been sampled (Denmark, Estonia, France, Latvia, the Netherlands, Poland and Switzerland) and European countries where a domestic lifecycle is established for E. granulosus (Estonia, France, Italy, Latvia, Poland, Portugal).

| Country (regions) | number of blueberries samples | *E. multilocularis* |  | *E. granulosus s.l.* |  | Other taeniid species |  |
| --- | --- | --- | --- | --- | --- | --- | --- |
|  |  | number of samples tested positive with *E. multilocularis* | proportions of positive cases (95% CI) | number of samples tested positive with *E. granulosus s.l.* | proportions of positive cases (95% CI) | number of samples tested positive with others taeniid species | proportions of positive cases (95% CI) with others taeniid species |
| Finland | 8 | 0 | 0% (0-36.9) | 0 | 0% (0-36.9) | 0 | 0% (0-36.9) |
| France | 31 | 1 | 3.2% (0.1-16.7) | 0 | 0% (0-11.2) | 0 | 0% (0-11.2) |
| Latvia | 30 | 4 | 13.3% (3.8-30.7) | 1 | 3.3% (0.1-17.2) | 0 | 0% (0-11.6) |
| Netherlands | 6 | 1 | 16.7% (0.4-64.1) | 0 | 0% (0-45.9) | 0 | 0% (0-45.9) |
| Poland | 3 | 0 | 0% (0-70.8) | 0 | 0% (0-70.8) | 0 | 0% (0-70.8) |
| Portugal | 15 | 0 | 0% (0-21.8) | 0 | 0% (0-21.8) | 0 | 0% (0-21.8) |
| Switzerland | 12 | 0 | 0% (0-26.5) | 0 | 0% (0-26.5) | 0 | 0% (0-26.5) |
| **SubTotal Europe\*** | 105 | 6 | 7.3% (2.7-15.2) | 1 | 1.3% (0.0-6.9) | 0 | 0% (0-3.5) |
| Pakistan | 25 | 14 | 56.0% (34.9-75.6) | 3 | 12.0% (2.5-31.2) | 1 *T. hydatigena*, 1 *Hydatigera* sp. | 8.0% (1.0-26.0) |
| **Total** | 130 | 20 | 15.4% (9.7-22.8) | 4 | 3.1% (0.8-7.7) | 1 *T. hydatigena*, 1 *Hydatigera* sp. | 1.5% (0.2-5.4) |

**Table 3:** Number of cases and proportions of detection of *E. multilocularis, E. granulosus sensu lato* and other taeniid species DNA in blueberries from the different countries. The 95% confidence intervals are indicated between brackets. The asterisk refers that the percentage value concerns only known endemic countries for *E. multilocularis*. *Proportions were calculated considering only those European countries for which areas endemic for *E. multilocularis* have been sampled (France, Latvia, the Netherlands, Poland and Switzerland) and European countries where a domestic lifecycle is established for *E. granulosus*(France, Latvia, Poland, Portugal).

| Country<br>(regions) | number<br>of<br>lettuces<br>analysed | *E. multilocularis* |  | *E. granulosus s.l.* |  | Other taeniid species |  |
| --- | --- | --- | --- | --- | --- | --- | --- |
|  |  | number of<br>samples tested<br>positive with<br>*E. multilocularis* | proportions of<br>positive cases<br>(95% CI) | number of<br>samples tested<br>positive with<br>*E. granulosus s.l.* | proportions of<br>positive cases<br>(95% CI) | number of samples<br>tested positive with<br>others taeniid<br>species | proportions of<br>positive cases<br>(95% CI) with<br>others taeniid<br>species |
| Denmark | 50 | 2 | 4.0% (0.5-13.7) | 0 | 0% (0-7.1) | 0 | 0% (0-7.1) |
| France | 226 | 3 | 1.3% (0.3-3.8) | 0 | 0% (0-1.6) | 6 *Hydatigera* sp. | 2.7% (1.0-5.7) |
| Germany<br>(Mecklenburg-Western<br>Pomerania) | 9 | 0 | 0% (0-33.6) | 0 | 0% (0-33.6) | 0 | 0% (0-33.6) |
| Germany<br>(Baden-Württemberg<br>region) | 63 | 0 | 0% (0-5.7) | 0 | 0% (0-5.7) | 1 *Hydatigera* sp. | 1.6% (0-8.5) |
| Italy (Abruzzo<br>region) | 80 | 0 | 0% (0-4.5) | 1 *E. granulosus*<br>s.s. | 1.3% (0-6.8) | 1 *Hydatigera* sp. | 1.3% (0-6.8) |
| Italy (Campania<br>region) | 46 | 0 | 0% (0-7.7) | 3 *E. granulosus*<br>s.s. | 6.5% (1.4-17.9) | 0 | 0% (0-7.7) |
| Italy (Sardinia<br>region) | 106 | 0 | 0% (0-3.4) | 4 *E. granulosus*<br>s.s. | 3.8% (1.0-9.4) | 3 *Hydatigera* sp.,<br>1 *T. multiceps* | 3.8% (1.0-9.4) |
| Latvia | 62 | 1 | 1.6% (0-8.7) | 1 *E. granulosus*<br>s.s. | 1.6% (0-8.7) | 1 *Hydatigera* sp. | 1.6% (0-8.7) |
| Netherlands | 6 | 0 | 0% (0-45.9) | 0 | 0% (0-45.9) | 0 | 0% (0-45.9) |
| Norway | 39 | 0 | 0% (0-9.0) | 0 | 0% (0-9.0) | 1 *T. krabbei/serialis* | 2.6% (0-13.5) |
| Poland | 74 | 0 | 0% (0-4.9) | 0 | 0% (0-4.9) | 0 | 0% (0-4.9) |
| Portugal | 101 | 0 | 0% (0-3.9) | 0 | 0% (0-3.9) | 0 | 0% (0-3.9) |
| Switzerland | 80 | 1 | 1.3% (0-6.8) | 0 | 0% (0-4.5) | 2 *Hydatigera* sp. | 2.5% (0.3-8.7) |
| **SubTotal**<br>**Europe\*** | 942 | 7 | 1.2%* (0.5-2.5) | 9 *E. granulosus*<br>s.s. | 1.3%*<br>(0.6-2.4) | 14 *Hydatigera* sp.,<br>2 *Taenia* sp. | 1.7% (1-2.7) |
| Pakistan | 100 | 2 | 2% (0.2-7.0) | 3 *E. granulosus*<br>s.s.,<br>1 *E. canadensis* | 4.0% (1.1-9.9) | 5 *T. saginata*,<br>3 *T. hydatigena*,<br>1 *Hydatigera* sp. | 9% (4.2-16.4) |
| Tunisia | 75 | 0 | 0% (0-4.8) | 9 *E. granulosus*<br>s.s. | 12.0% (5.6-21.6) | 3 *T. hydatigena* | 4% (0.8-11.2) |
| **Total** | 1,117 | 9 | 0.8% (0.4-1.5) | 21 *E. granulosus*<br>s.s.,<br>1 *E. canadensis* | 2.0% (1.2-3.0) | 15 *Hydatigera* sp., 13<br>*Taenia* sp. | 2.5% (0.8-11.2) |

The species identification of *E. granulosus s.l*. DNA from berries was limited to *E. granulosus s.s*. The overall contamination levels in this study were 5.3% (16/300) in strawberries and 3.1% (4/130) in blueberries (Tables 2 and 3). *Echinococcus granulosus s.l*. DNA was only detected in countries where the domestic life cycle of the parasite is known to be present, such as Italy, Latvia, Pakistan, Poland, Portugal, and Tunisia. In Europe, the DNA was detected in 1.1% (3/284) of strawberries and 1% (1/105) of blueberries. In European areas from the Mediterranean basin (Portugal and Sardinia) and Eastern Europe (Estonia, Latvia, and Poland) which are endemic for CE, the contamination rate of *E. granulosus s.s*. in strawberries (3/145) and blueberries (1/48) was 2.1% for both. The level of contamination varied from 1.8% (1/55) in Sardinia, to 5.9% (1/17) in Portugal and was highest in Poland with 7.7% (1/13) of strawberries testing positive for *E. granulosus s.s.* DNA. *Echinococcus granulosus s.s.* DNA in blueberries was detected in 3.3% (1/30) of samples from Latvia. Higher contamination levels of *E. granulosus s.s.* DNA were found outside of Europe, with 12.0% (3/25) of in blueberries from Pakistan and 81.3% (13/16) of strawberries from Tunisia. No *E. granulosus s.s*. DNA was detected in other types of berries (Supplementary Table 3).

Other taeniid species, including *Hydatigera* sp. and *T. hydatigena*, were detected in strawberries and blueberries during this study. Additionally, *Mesocestoides melesi* was detected in blueberries from Latvia. The overall detection level of other taeniid species DNA was 1.3% (4/300) and 1.5% (2/130) in strawberries and blueberries, respectively (Tables 2 and 3). In Europe, only *Hydatigera* sp. was detected in strawberries at 1.1% (3/284). In Tunisia, *T. hydatigena* was detected in 6.3% (1/16) of strawberries. Contamination of blueberries with both *Hydatigera* sp. and *T. hydatigena* was observed at 8% (1/25; 1/25) in Pakistan. No taeniid species DNA others than *Echinococcus* spp. were identified from other berries (Supplementary Table 3).

### 3.4. Differences in contamination according to type of samples and supplying sources

As regards the supply sources of samples, the majority of lettuces (76.2%; 851/1,117) was purchased from local markets, 19% (212/1,117) from local gardens and only 4.8% (54/1,117) from supermarkets. *Echinococcus multilocularis* and *E. granulosus s.l.* DNAs were detected in lettuces from all supply sources except *E. granulosus s.l.*, which was not detected in supermarket lettuces. Strawberries were obtained from local markets (85%; 255/300), kitchen gardens (10%; 30/300), supermarkets (4.7%; 14/300) and forest or field (0.3%; 1/300). The blueberries were mainly collected from forests (58.5%; 76/130), 35.4% (46/130) from local markets and 6.2% (8/130) from private gardens. *Echinococcus multilocularis* and *E. granulosus s.s*. DNAs were only found in blueberries from private kitchen gardens and local markets, whilst both parasite DNA was found in strawberries from all supply sources. Regarding the different supply sources of lettuces in Europe (private kitchen gardens, local markets, supermarkets), no significant difference were observed in the contamination by *E. multilocularis* (p=0.057) or by *E. granulosus s.l.* (p=0.157). The same observation was seen with strawberries (p=0.404 for *E. multilocularis* and p= 0.307 for *E. granulosus s.l.*) and blueberries (p=1 for *E. multilocularis* and p= 0.294 for *E. granulosus s.l.*). No significant difference (p=0.776) was observed between the proportion of *E. granulosus s.s*. DNA contaminating lettuces, strawberries, and blueberries in European endemic areas. However, in European endemic areas, a significantly higher proportion of *E. multilocularis* contamination was found in strawberries and blueberries compared to lettuces (p=0.0002). Additionally, a significantly higher proportion of contamination by *E. granulosus s.s*. was observed in strawberries compared to lettuces in Tunisia (p<0.00001). A significantly higher proportion of *E. multilocularis* contamination was observed in blueberries compared to lettuces (p<0.00001) from Pakistan. Despite having higher proportions of *E. granulosus s.l*. contamination in blueberries than in lettuces in Pakistan, significance level was not reached (p=0.142).

## 4. Discussion

Due to the long incubation period of *E. multilocularis* and *E. granulosus s.l*., human CE and AE have been historically considered foodborne diseases, even in the absence of strong evidence of food transmission. Therefore, the acquisition of data on food contamination with *Echinococcus* spp. is crucial to assess the food-borne transmission risk. This study focuses on lettuce and berry, as these products are food items commonly consumed raw, sometimes unwashed, that can easily become contaminated by parasite eggs on the ground and have been considered as a potential source of human infection. In this study, the food items were not washed before being processed in order to provide data on contamination, independently from the washing that should be realized before consumption. To detect the presence of *Echinococcus* spp. DNA in food samples, a sensitive detection method was necessary. Guggisberg et al. (2020) published a one-way sequential sieving method for lettuces, which was followed by a multiplex-PCR designed to distinguish *E. multilocularis, E. granulosus s.l.* and others taeniid species (Trachsel et al., 2007). In these two multicenter studies, a real-time PCR detection approach was chosen for *E. multilocularis* and *E. granulosus s.l.* (Knapp et al., 2016; Maksimov et al., 2020). The limit of detection in the conditions laboratory of the multicenter studies was evaluated for *E. multilocularis* in lettuces and also for the first time in berries. Only a minor adjustment to the method was required to remove the eggs without damaging the berries, which could have complicated further filtration steps. In this study, it is assumed that the detected parasite DNA originates from eggs due to the mesh size used to filter the samples. The method demonstrated high sensitivity, detecting up to three eggs in lettuce and berry samples, and even one or two eggs in very high proportions of spiked samples. The sampling of lettuce and berries was designed to minimize the number of samples originating from supermarkets, as any washing or other treatment at manufacturing level would have an impact on the probability to detect parasite DNA. Among the limits of this multicenter study, we should acknowledge possible sampling bias due to: i) the number and varieties of samples collected from one country to another (especially for berries), and ii) geographical origin of samples, generally limited from few areas of the country. Additionally, the percentage of detections gave an overview of the contamination of samples from an area in each country but may not be considered as an absolute value for the country as a whole.

The country-based proportions of lettuce and berry samples contaminated by *E. multilocularis* and *E. granulosus s.l.,* observed in this multicenter study were consistent with the known presence of both parasites in these countries. For instance, *E. granulosus s.l.* was not detected in lettuces and berries from countries where the domestic lifecycle of the parasite is not present, such as Denmark, the Netherlands, Germany, and Finland or where the parasite is present at low endemicity such as France (Deplazes et al., 2017). *Echinococcus granulosus s.s.* was predominantly detected in endemic countries where a dog-sheep life cycle occurs, such as Italy, Tunisia and Pakistan. This parasite species was detected in all three endemic Italian regions sampled, confirming a high level of environmental contamination. This is consistent with the strong transmission dynamic between dogs, sheep and humans in the country (Casulli, Abela-Ridder, et al., 2023; Cringoli et al., 2007; Piseddu et al., 2017). In contrast, the parasite was not detected in lettuces from Portugal, where the parasite is present, but at low prevalence in animals and low incidence in humans, respectively (Casulli, Abela-Ridder, et al., 2023; EFSA, 2022). However, it should be noted that sampling was only conducted in two local markets from two different villages in Portugal. *Echinococcus granulosus s.s*. was also detected in berries from Latvia and Poland and in one lettuce from Latvia, both eastern European countries where the domestic lifecycle of the parasite is present. In this study, the highest contamination level for *E. granulosus s.s*. was detected from Tunisia, in both lettuces and strawberries. The strawberries were all grown in Cap Bon, located in the northeast of Tunisia, which is the only region in the country with a milder climate suitable for strawberry production. For lettuce, contaminated samples were collected from different regions of the country, including 16 samples in Cap Bon, where three (18.8%) samples tested positive to *E. granulosus s.s.* This may indicate a high level of environmental contamination in the country, which is consistent with the high prevalence reported in both humans and animals in Tunisia (M’Rad et al., 2020). The level of contamination in lettuces from Pakistan was similar to that of lettuces from Italy, but blueberries had a higher contamination rate of 12%. These values from Pakistan and Tunisia are both higher than those from Europe. It should be noted that the only *E. canadensis* (G7) specimen has been identified in one lettuce from Pakistan. These results emphasize the need to investigate the environmental contamination and the agricultural practices to produce specific vegetables and berries in different continents to better understand the potential risk associated to foodborne transmission of human CE.

*Echinococcus multilocularis* DNA was detected in food items from seven (Denmark Estonia, France, Germany, Latvia, the Netherlands and Switzerland) of the eight European endemic countries (EFSA, 2022) sampled in this study, as well as in Pakistan, where its presence has also been previously reported (Khan et al., 2020; Khan et al., 2021). The parasite is currently considered as absent from Finland, Norway (except for Svalbard Archipelago) and Portugal but also from Tunisia. In Italy, *E. multilocularis* is only present in the northern part of the country.

The current northern distribution of *E. multilocularis* in Italy, represented by Liguria region bordering France and the Trentino-Alto Adige region bordering Austria (Casulli et al., 2005; Massolo et al., 2018), does not overlap with the central-southern regions sampled in this study (Abruzzo, Campania and Sardinia). For this reason, the absence of *E. multilocularis* detection in lettuces and strawberries from Italy was not surprising. In endemic countries of Europe, the prevalence of *E. multilocularis* in red foxes, which can be used as an indirect measure of environmental contamination by eggs, varies significantly from one country to another (Oksanen et al., 2016). Therefore, the absence of detection of *E. multilocularis* in both lettuces and berries from Poland may seem surprising in comparison to results from other endemic countries. Indeed, not only because of the high prevalence reported in red foxes (Karamon et al., 2014), but also due to the high percentage of the reported contamination in forest fruits, vegetables and mushrooms collected from both high and low Polish endemic areas from previous studies (Lass et al., 2015b, 2017). In our study, lettuces from Poland were mainly purchased in local markets, while strawberries were collected from kitchen gardens and blueberries from the forest. This sampling was similar to that carried out in the two previous studies from Poland. However, some concerns were raised on the methodology and potential cross-contamination in those two studies which could draw the results into question (Alvarez Rojas et al., 2018; Lass et al., 2016; Robertson et al., 2016; Torgerson, 2016). In contrast to Poland, 4% of lettuce samples (n=50) and 1.8% of strawberry samples (n=56) from Denmark were found to be contaminated by *E. multilocularis*. Interestingly, such percentages has not been found from an area where high local prevalence of *E. multilocularis* in foxes have been previously described (Petersen et al., 2018), reflecting the patchy distribution of *E. multilocularis* at a local scale. This highlights the difficulty in drawing definitive conclusions regarding the different levels of food contamination between countries, especially due to the relatively low number of samples collected in each area. To resume, similar values of around 1% *E. multilocularis* DNA were obtained in the three European countries (France, Latvia, and Switzerland), where the parasite was identified in lettuces. These findings also match a prior study from Switzerland, whereby a similar proportion of contaminated 1.3% lettuces (n=157) was found using the same washing and sieving method (Guggisberg et al., 2020). Based on this data, it can be estimated from this study that overall 1% of lettuces sampled in endemic areas of European countries are contaminated by *E. multilocularis*.

Among the few previous studies on the detection of *E. multilocularis* in berries, there was no detection in blueberry samples from Estonia (n=21) and Finland (n=21) (Malkamäki et al., 2019). *E. multilocularis* DNA was identified in 20% of raspberry samples from plantations in Poland (Lass et al., 2015a). In contrast, our study detected the parasite in berries from all six European endemic countries, with values frequently exceeding 12%, however none were found in the samples from Poland. Due to the limited sample size of berries tested for *E. multilocularis*, the absence of detection of the parasite in the 30 berry samples from Poland is perhaps not surprising. Nor was *E. multilocularis* DNA found in blueberries (n=12) and strawberries (n=11) from the highly endemic country of Switzerland, but one of the six additional raspberry samples tested positive. In this study, the highest contamination level by *E. multilocularis* was obtained from blueberries in Pakistan and raises questions about the endemic level in the country and on the level of hygiene during production. Contaminated samples were collected from local markets, vegetable gardens, and from the wild from all the five different cities of production sampled in Pakistan. The incidence of AE in Pakistan is considered to be very low, with only a few human cases reported and a low occurrence in 3 out of 68 (4.4%) red fox faeces examined from the north of the country (Borhani et al., 2021; Khan et al., 2021). Additional epidemiological data on *E. multilocularis* is required from Pakistan for humans, animals, and foodstuffs, such as berries and vegetables, to obtain a more accurate evaluation of the potential risk of foodborne transmission. Furthermore, the potential contribution of dogs to the lifecycle of *E. multilocularis* must be explored in Pakistan.

In addition to *Echinococcus* spp., DNA from several other taeniid species were identified, confirming the ability of taeniid eggs to be mechanically transferred on food. In Europe, besides *Echinococcus* spp., the most frequently identified taeniid species (21 out of the 23 taeniid DNA detections) in this study was from the genus *Hydatigera*, resulting from cat defecation. Our finding is consistent with the identification of kitchen gardens as hotspots for both fox and cat defecation in northeastern France (Bastien et al., 2018). Since only a few cats may sporadically excrete high numbers of *E. multilocularis* eggs (Umhang et al., 2022), this species is considered to have a very low biotic potential and therefore does not significantly contribute to overall *E. multilocularis* environmental contamination (Hegglin and Deplazes, 2013; Kapel et al., 2006). Additionally, the *E. multilocularis* eggs obtained from cats are generally considered to be non-infectious, which strongly limits the role of cats in foodborne transmission of AE. In a lettuce sample from Norway, it was impossible to differentiate between *T. serialis* and *T. krabbei* based on the partial 12S gene sequence. Both species are present in the country and are associated with intermediate hosts in wildlife (i.e. rabbits and cervids, respectively) and canids (dogs, foxes or wolves) as the definitive hosts. *Taenia hydatigena* was detected in both lettuces and berries in Tunisia and Pakistan. This maybe expected according to the detection in high proportions of *E. granulosus s.s*. in both countries as these species share the same lifecycle between dogs and livestock. The detection of *T. saginata* in Pakistan is associated with environmental contamination by human feces, as humans are the only definitive host. The presence of *T. multiceps* in Sardinia is consistent with previous reports on the presence of this parasite on this island (Scala et al., 2007). This seems to confirms the hypothesis that dogs, which consume raw sheep offal and live in proximity to lettuce and strawberry fields may be a source of *E. granulosus s.s.* eggs contamination. It is worth noting that *Taenia* worms have a much higher egg production capacity than *Echinococcus* worms (Alvarez Rojas et al., 2018). Nevertheless, the rate of DNA detection from *Echinococcus* was found to be globally comparable to that of other taeniid in lettuces and higher in berries, according to this study. In comparison, this was not the case in Guggisberg et al. (2020) study which detected 10 lettuce samples with DNA of others taeniid species versus two with *E. multilocularis* (n=157) and Federer et al. (2015) found only two *E. granulosus* positives mixed vegetables and fruits samples versus 23 positive for others taeniid species among 141 samples. The use of real-time PCR assays for detection of both *E. multilocularis* and *E. granulosus s.l*. in the present study, compared to classical PCR in the other, may support a relatively higher sensitivity of *Echinococcus* species with real-time PCR assays. However, the diversity of *Taenia* species identified was lower than that in the two Swiss studies, despite using the same classical PCR method. Notably, *Taenia crassiceps* and *Taenia polyacantha*, which are typically found in foxes, were not detected in our study.

The highest *E. multilocularis* contamination rate of berries compared to lettuces may be due to a difference in the shapes and surface porosity of these vegetables with different egg retention on the different surfaces. Malkamäki et al. (2019) reported a difference in taeniid egg contamination between two types of berries. Since a difference of contamination was not observed for *E. granulosus s.s*. in Europe, it can be supposed that red foxes are more likely to be close to berries than dogs. In Tunisia, the strawberry samples were exclusively obtained from a region which have a milder climate, potentially suggesting a better survival of eggs even if comparable proportions of lettuces contaminated by *E. granulosus s.s.* were obtained from there. The situation in Pakistan is interesting because it is endemic for both CE and AE, with dogs assumed to be strongly involved in the dispersion of eggs of both parasite species. Once again, the difference in contamination between lettuces and blueberries was observed only for *E. multilocularis*. It cannot be assumed that the proportion of parasitic contamination detected in the different food items can be linearly related to the risk of human infection. This is because many factors were not considered, such as the relative quantities of lettuces and berries eaten by consumers or the different exposure of people due to different cultural habits, like prewashing of vegetables and fruits or the proportion of food imported from endemic countries, to obtain a more accurate risk assessment of human foodborne contamination by *Echinococcus* species.

The global contamination rate of taeniid eggs in lettuces (5.3%) and berries (13.3%) suggests that the transfer of taeniid eggs from carnivore feces to food is not uncommon. However, although we assume that eggs are the source of the DNA detected in this study, the viability of such eggs is unknown. To obtain direct evidence and an accurate risk assessment of foodborne human infection by *E. multilocularis* and *E. granulosus s.l*., it is necessary to firstly confirm the viability of *Echinococcus* eggs in food. Currently, in the absence of sensitive in vitro methods, the most reliable viability test requires the experimental infection of small rodents with *Echinococcus* eggs. The subcutaneous method involving mice has been revealed as the most sensitive inoculation method, rather than intraperitoneal or peroral infection (Alvarez Rojas et al., 2018; Federer et al., 2015).

The relative contribution of foodborne infection as compared to other sources (e.g. direct contact with dogs, water, or other environmental sources) is likely to differ between continents for both AE and CE (Torgerson et al., 2020). The detection of *E. multilocularis* and *E. granulosus s.l*. in lettuces and berries in this study suggests a potential risk of foodborne human infection. The relative contribution of this risk remains to be determined. In the future it would be also important to prioritize studies on novel viability tests for the eggs isolated in food matrices as well as other studies about parasitic DNA detection in food. Moreover, further studies are necessary to combine the detection of *Echinococcus* eggs in food with other environmental matrices from various continents. This will help to assess different epidemiological contexts and social habits, leading to a better understanding of human infections by *Echinococcus* eggs.

## Acknowledgments

The authors thanks Justine Pellegrini and Vanessa Bastid for technical contribution regarding analyses of blueberries from Pakistan. This work was supported by funding from the European Union’s Horizon 2020 Research and Innovation programme under grant agreement number 773830: One Health European Joint Programme (MEME project; https://onehealthejp.eu/jrp-meme/). This work was also partially supported by the European Commission’s Single Market Programme (SMP Food) under the grant agreement no. 101144113: “Work programme 2023-2024 of EU European Reference Laboratory for the Parasites (EURLP)”. This work was also supported by research funding (grant PRG1209) from the Estonian Ministry of Education and Research (to US). For the purpose of Open Access, the author has applied a CC-BY public copyright license to any Author Accepted Manuscript (AAM) version arising from this submission.

