## Supplementary table 1 for "Detection of *Echinococcus* spp. and other taeniid species in lettuces and berries: two international multicenter studies from the MEmE project"

| **Supplementary table 2**: Number of cases and proportion of detection of *Echinococcus multilocularis*, *Echinococcus granulosus sensu lato*and others taeniid species DNA in others vegetables than lettuces from the different countries. The 95% confidence intervals are indicated between brackets. *Proportions were calculated considering only those European countries for which areas endemic for *E. multilocularis* have been sampled (Germany, Latvia and Switzerland) and European countries where a domestic lifecycle is established for *E. granulosus* (Italy and Latvia). | | | | | | | | | | |
| --- | --- | --- | --- | --- | --- | --- | --- | --- | --- | --- |
|  |  |  | ***E. multilocularis*** | |  | ***E. granulosus s.l.*** | |  | **Others taeniid species** | |
| **Country (regions)** | **number of vegetables others than lettuces analysed** | **type of other vegetables analysed** | **number of samples tested positive with *E. multilocularis*** | **proportions of positive cases (95% CI)** |  | **number of samples tested positive with  *E. granulosus s.l.*** | **proportions of positive cases (95% CI)** |  | **number of samples tested positive with others taeniid species** | **proportions of positive cases (95% CI) with others taeniid species** |
| Germany (Mecklenburg-Western Pomerania) | 11 | 2 carrot leaves, 6 chard, 1 onion, 1 parsley romarin, 1 thyme | 1 (chard) | 9.1%  (0.2-41.3) |  | 0 | 0%  (0-28.5) |  | 0 | 0%  (0-28.5) |
| Italy (Campania) | 4 | 4 Chicory endive | 0 | 0% (0-60.2) |  | 0 | 0% (0-60.2) |  | 0 | 0% (0-60.2) |
| Italy (Sardinia) | 1 | 1 chard | 0 | 0% (0-97.5) |  | 0 | 0% (0-97.5) |  | 0 | 0% (0-97.5) |
| Latvia | 38 | 4 Basil, 2 beet leaves, 1 coriander, 1 kale, 14 parsley, 2 celery leaves, 5 sorrel, 9 spinach | 0 | 0% (0-9.3) |  | 0 | 0% (0-9.3) |  | 3 *Hydatigera* sp. from 2 sorrel  and 1 basil | 7.9% (1.7-21.4) |
| Norway | 11 | 11 chard | 0 | 0% (0-28.5) |  | 0 | 0% (0-28.5) |  | 1 *Hydatigera* sp. | 9.1% (0.2-41.3) |
| Switzerland | 6 | 4 red chicory, 2 stem lettuces | 0 | 0% (0-45.9) |  | 0 | 0% (0-45.9) |  | 0 | 0% (0-45.9) |
| **Total*** | 71 |  | 1 | 1.8%* (0-9.7) |  | 0 | 0%* (0-8.2) |  | 4 *Hydatigera*sp. | 5.6% (1.6-13.8) |
