## Supplementary table 2 for "Detection of *Echinococcus* spp. and other taeniid species in lettuces and berries: two international multicenter studies from the MEmE project"

| **Supplementary table 3**: Number of cases and proportions of detection of *E. multilocularis*, *E. granulosus sensu lato* and other taeniid species DNA in berries others than strawberries and blueberries from the different countries. The 95% confidence intervals are indicated between brackets. *Proportions were calculated considering only those European countries for which areas endemic for *E. multilocularis* have been sampled (Netherlands, Poland and Switzerland) and European countries where a domestic lifecycle is established for *E. granulosus* (Poland and Portugal). | | | | | | | | | | | |
| --- | --- | --- | --- | --- | --- | --- | --- | --- | --- | --- | --- |
|  |  |  |  | ***E. multilocularis*** | |  | ***E. granulosus s.l.*** | |  | **Other taeniid species** | |
| **Country** | **number of berries samples analysed** | **details of the others berries samples** |  | **number of samples tested positive with *E. multilocularis*** | **proportions of positive cases (95% CI)** |  | **number of samples tested positive with  *E. granulosus s.l.*** | **proportions of positive cases (95% CI)** |  | **number of samples tested positive with others taeniid species** | **proportions of positive cases (95% CI) with others taeniid species** |
| Finland | 7 | 2 raspberries, 3 red lingonberry, 2 blackcurrant |  | 0 | 0% (0-41.0) |  | 0 | 0% (0-41.0) |  | 0 | 0% (0-41.0) |
| Netherlands | 11 | 3 blackberries, 4 red currants, 1 white currants, 3 raspberries |  | 0 | 0% (0-28.5) |  | 0 | 0% (0-28.5) |  | 0 | 0% (0-28.5) |
| Poland | 14 | 4 blackberries, 4 blueberrry bushes, 5 raspberries, 1 Saskatoon berry |  | 0 | 0% (0-23.2) |  | 0 | 0% (0-23.2) |  | 0 | 0% (0-23.2) |
| Portugal | 5 | 5 Raspberries |  | 0 | 0% (0-52.2) |  | 0 | 0% (0-52.2) |  | 0 | 0% (0-52.2) |
| Switzerland | 13 | 6 blackberries, 1 red currant, 6 raspberries |  | 1 | 7.7% (0.2-36.0) |  | 0 | 0% (0-24.7) |  | 0 | 0% (0-24.7) |
| **Total*** | 50 | |  | 1 | 2.6%* (0-13.8) |  | 0 | 0%* (0-7.0) |  | 0 | 0% (0-17.6) |
