## Supplementary table 3 for "Detection of *Echinococcus* spp. and other taeniid species in lettuces and berries: two international multicenter studies from the MEmE project"

| **Supplementary Table 1:** Estimation of the limit of detection of the washing and sequential sieving method combined with real-time PCR detection of *E. multilocularis* DNA from lettuces (300g) and strawberries (200g) samples spiked with ten to one *Echinococcus multilocularis* eggs. The recovery results are presented by the proportion of tested positives out of the number of replicates tested and percentages of detection. | | | | | | |
| --- | --- | --- | --- | --- | --- | --- |
|  | Number of eggs spiked | Number of positive lettuces out of tested | % |  | Number of positive strawberries out of tested | % |
|  | 10 | 10/10 | 100 |  | 3/3 | 100 |
|  | 5 | 5/5 | 100 |  | 2/2 | 100 |
|  | 3 | 23/24 | 95.8 |  | 8/8 | 100 |
|  | 2 | 21/24 | 87.5 |  | 7/8 | 87.5 |
|  | 1 | 12/24 | 50 |  | 7/8 | 87.5 |
